# Immersive bottle effect: Influence of virtual object simulation on modulation of corticospinal excitability by motor imagery

**DOI:** 10.64898/2026.06.02.729504

**Authors:** Clémence Bonnet, Mélina Behava, Stéphane Argon, Sidney Grosprêtre

## Abstract

Motor imagery is a cognitive process that engages the motor system and facilitates motor learning, recovery, and performance. Its effectiveness relies on the activation of corticospinal pathways, which can be modulated by action observation. This study investigated whether observing graspable objects facilitates corticospinal excitability during motor imagery, and whether immersive virtual reality amplifies this effect. Twelve healthy adults of either sex completed a single session involving six conditions: rest, object observation on a screen or in virtual reality, motor imagery alone, and combined observation with motor imagery. Transcranial magnetic stimulation was used to record motor-evoked potentials from hand and forearm muscles, and fatigue and imagery quality were quantified. Results showed that observing a manipulable object during motor imagery significantly increased corticospinal excitability. This facilitation was strongest when the object was presented in a virtual environment. The effect was muscle-specific, targeting the agonist hand muscle of the imagined action (Abductor Pollicis Brevis), and was more pronounced at the higher stimulation intensity. Fatigue and imagery quality were similar across conditions. These findings indicate that object affordances can prime motor circuits and enhance motor imagery-induced neural activation, with immersive environments further reinforcing this effect. This is the first study demonstrating that combining motor imagery with virtual object observation maximizes corticospinal excitability. This approach may represent a promising tool for rehabilitation and sports training. Further studies should identify the optimal parameters and the neurophysiological markers of cortical modulation for this combination.

**NEW & NOTEWORTHY:** Using transcranial magnetic stimulation with immersive virtual reality, we showed for the first time that observing a virtual object associated with the imagined movement increased corticospinal excitability to a greater extent than motor imagery performed alone.

## INTRODUCTION

Provide a brief overview of the scope and relevance of the study, especially regarding previous advancements in related fields.

### Motor imagery activates the nervous system

Motor imagery (MI) is a cognitive process involving the mental simulation of a motor action in the absence of overt movement (1,2). MI recruits neural networks that partially overlap with those involved in motor preparation and execution, including supplementary motor area, premotor and primary cortices, parietal cortex, cerebellum, basal ganglia, and thalamus (3,5–9). This shared neural recruitment is associated with increased corticospinal excitability (10–12) and modulation of the spinal network (13,14). The overlap between the neural substrates of MI and motor execution is thought to contribute to the effects of MI on motor learning and performance, supporting its application in rehabilitation (7,15–17), sports performance (18–20), and musical practice (21,22). Whereas visual MI (i.e., imagining seeing yourself moving) primarily recruits occipital areas, kinesthetic MI (i.e., imagining feelings of movement) elicits stronger activation in motor-related areas, (7,23,24), although lower than actual movement execution (5). Bouguetoch et al. (2020) suggested that MI could be an intermediate level of corticospinal activation between a slight voluntary contraction and rest (25). Similarly, a meta-analysis demonstrated that MI practice improves performance, although to a lesser extent than physical practice (26). Overall, these findings highlight the need to develop new strategies that further enhance the effectiveness of MI.

### Enhanced activation during combined motor imagery and action observation

Beyond MI alone, action observation (AO) also engages the motor system by activating the mirror neuronal network and increases corticospinal excitability (27–29). Over the past few years, many studies have sought to enhance the effectiveness of MI by combining it with AO. This combined approach, in which individuals imagine an action while observing it (AO+MI), has been shown to be more effective than MI alone (30–33). Specifically, AO+MI increased corticospinal excitability (34,35) and activity in motor-related brain areas (36), compared to either MI or AO alone. Such findings are promising for patients (37,38) and athletes (39–41). Interestingly, some recent studies reported enhanced MI while observing the imagined action in an immersive virtual reality (VR) headset (42–44). VR is a technology that recreates realistic environments and provides artificial sensation to the users in a three-dimensional space. A key feature of VR is the feeling of “being there” in the virtual environment, commonly called “the sense of presence”, that has been linked to neurophysiological markers (45,46). Despite consistent evidence showing increased activity in motor brain regions during AO+MI, the combination can sometimes hinder the imagery process, making it harder for the individual to internalize the gesture. A common issue arises from the perspective mismatch: while the observed movement is performed by another person in most cases, participants are usually instructed to imagine themselves executing the same movement. Generating a vivid kinesthetic imagery under these conditions can be challenging (47). Indeed, a third-person video must need to be mentally transformed into a first-person perspective to align the observed action with the internal simulation of movement parameters (e.g., timing, amplitude, sensations). There can also be a temporal mismatch between the natural and spontaneous pace of an individual to mentally perform an action and the same action observed through a third person perspective. Another important consideration concerns individuals with advanced age or neurological impairments. For theses populations, the AO+MI combination may be particularly demanding (30). This highlights the need to refine action observation protocols to maximize their facilitative role in MI rather than interfere with it, for instance by displaying only the environment in which the action takes place or the objects involved.

### Affordances and motor system activation

It is now admitted that observing graspable objects such as a cup or a teapot automatically engages the motor cerebral network (48). This activation is due to affordances, defined as the action possibilities offered by an object for interaction (49). In his theory of action affordance, Gibson (1979) stated that objects offer possibilities for interactions with the environment (50). In their cognitive theory, Tucker & Ellis (1998) postulated that objects (e.g., a bottle) would prime motor actions typically associated with them (e.g., grasping; (51). Indeed, reaction times decreased in an object-grip compatibility task when a small object is associated with a precision grip, and a large object is associated with a power grip (52,53). This compatibility showed that seeing a grasping object triggers a visuomotor priming that activates corresponding motor pattern (54,55). Supporting these behavioral findings, PET and fMRI studies reported activation in motor-associated brain areas during the perception of graspable objects (48,56,57). More recently, Schulz et al. (2018) showed that MI of a grasping movement involving a familiar object induced stronger recruitment of motor-related regions compared to a geometrical shape, highlighting the contribution of visuo-spatial cognition. Despite the clear link between affordances and action, the influence of object observation (OO) during MI remains poorly understood. It is particularly unclear whether the presence of affordances modulates corticospinal excitability during combined OO+MI. Furthermore, now technological advances allow displaying objects and environments in a more realistic and immersive way, i.e. by using VR instead of a simple 2D screen. Yet, the added value of VR compared to traditional screen-based presentation remains to be investigated.

The aim of the present study was twofold. First, we investigated whether affordances modulate corticospinal excitability during MI of grasping involving a familiar everyday object (i.e., a water bottle). Using transcranial magnetic stimulation (TMS) to measure motor evoked potentials of hand and forearm muscles, we hypothesized an increased corticospinal excitability in the combined OO+MI condition compared to either OO or MI alone, reflecting increased engagement of motor-related neural networks. Second, we examined the role of the display medium—screen or VR headset—in presenting the object, to assess whether an immersive environment amplifies the presumed effect of combined OO+MI. We hypothesized more pronounced effects of affordances on corticospinal excitability when presented in VR.

## MATERIALS AND METHODS

### Ethical approval

All participants were informed of the experimental protocol and potential discomfort, and they provided written informed consent prior to the session. The study was approved by the national French Ethical Committee [Institutional Review Board: Comité de Protection des Personnes (CPP)-Est-4, number 18/48], and research procedure complied with the *Declaration of Helsinki* (1964, amended).

### Participants

An a priori power analysis was conducted to determine the required sample size, based on the effect of combined AO+MI on corticospinal excitability reported in a previous study, as assessed by motor-evoked potentials amplitude recorded from the first dorsal interosseous during imagined right index finger abduction (38). Based on their reported significant muscle × video interaction effect [*F*(5,90) = 4.32; *p* = 0.001; 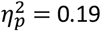], corresponding to a standardized effect size of *f* = 0.484, an a priori power analysis for a repeated-measures ANOVA (*α* = 0.05, power = 0.95) indicated a minimum required sample size of *N* = 6 (G*Power ; Faul et al., 2007). Our current proposed sample size of *N* = 12 exceeded this estimate.

The experimental group included 12 healthy young adults (4 women, 8 men; age: 27.25 ± 6.9 years; height: 1.72 ± 0.07 m; weight: 70.45 ± 12.96 kg, mean ± *SD*). All participants were right-handed, as confirmed by the Edinburgh Handedness Inventory (mean score: 0.84; Oldfield, 1971). Prior to participating in the experiment, all participants reported no neurological or physical disorders. Those with contraindications to TMS or nerve stimulation were excluded.

### Experimental design

Participants completed a single 2.5-hour experimental session. They were seated comfortably in a chair, with their hands and arms resting in a relaxed position on their thighs (Figure 1.A). The visual stimulus was presented either on a 42.51-inch DELL monitor (3840 × 2160 pixels; 60 Hz) positioned 1.70 m away, or via an Oculus Quest 2 VR headset (Meta Platforms, Irvine, US). Both devices were connected to a DELL computer equipped with an NVIDIA T600 Laptop GPU to ensure stable and high-quality visual stimulus presentation. Participants were instructed to remain as still as possible and to avoid voluntary movements throughout the session.

**Figure 1.**
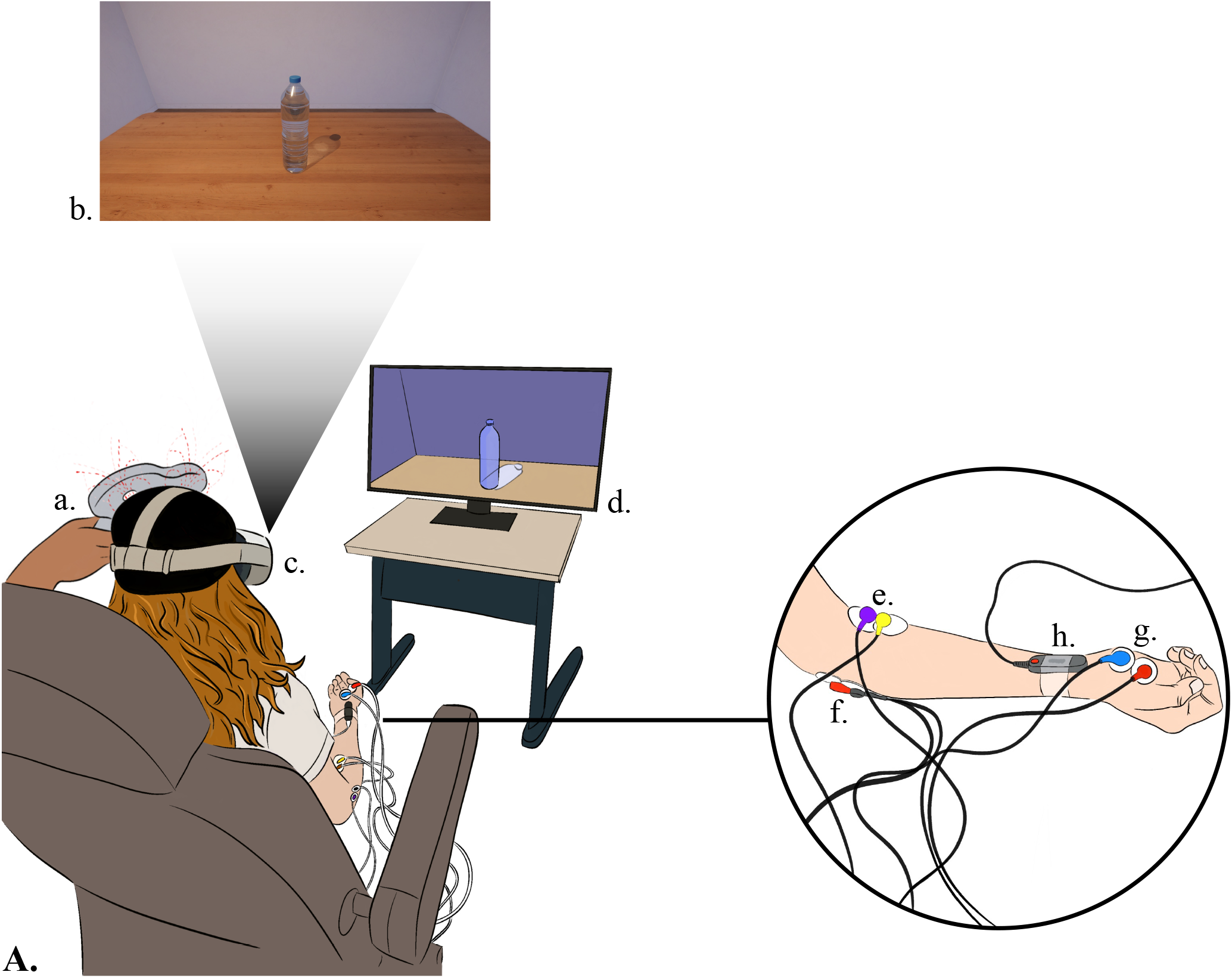
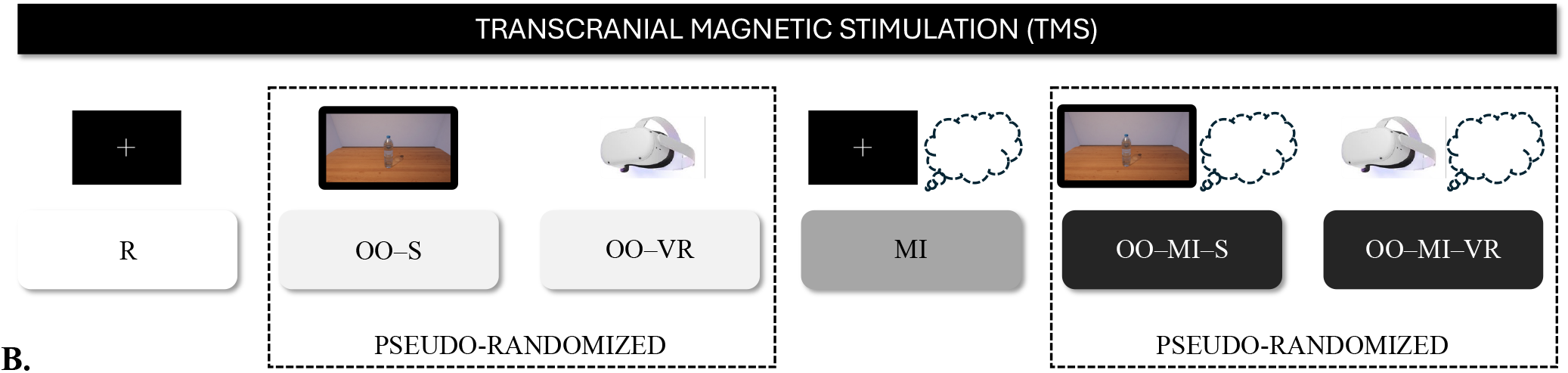
Experimental protocol. Figure 1.A – Experimental setup. **a**. Transcranial Magnetic Stimulation (TMS) figure eight-coil over the left motor cortex; **b**. Virtual environment used in the screen and the virtual reality headset; **c**. Virtual reality headset; **d**. Screen; **e**. Palmaris Longus; **f**. Extensor Carpi Radialis; **g**. Abductor Pollicis Brevis; **h**. Stimulator of median nerve. The distance between the participant and the screen was fixed at 1.70 m. **Figure 1.B – Experimental design. R**: Rest; **OO–S**: Object Observation Screen; **OO–VR**: Object Observation Virtual Reality; **MI**: Motor Imagery; **OO–MI–S**: Object Observation Motor Imagery Screen; **OO–MI–VR**: Object Observation Motor Virtual Reality. The R condition was always performed first, followed by the two OO conditions. Then the MI condition was performed, followed by the two combined OO–MI conditions. Each condition comprised 10 trials during which a transcranial magnetic stimulation was applied.

The procedure is illustrated in Figure 1.B After completing the questionnaires, participants took part in six different conditions (four experimental and two control; Table 1), involving each 12 trials, with a 2-minute interval between conditions. During ten trials for each condition, motor-evoked potentials (MEPs) elicited by TMS were evoked from the target muscle Abductor Pollicis Brevis (APB). MEPs were also measured from the Palmaris Longus (PL) and the Extensor Carpi Radialis (ECR). During two trials for each condition, M-waves were elicited by means of peripheral nerve stimulation.

**Table 1.** Description of experimental conditions.

| Acronym | Name | Task | Definition |
| --- | --- | --- | --- |
| R | Rest | Rest | Fixation on a cross at the center of the screen |
| OO-S | Object Observation Screen | Observation | Observing a full water bottle on the screen |
| OO-VR | Object Observation Virtual Reality | Observation | Observing a full water bottle through the VR headset |
| MI | Motor Imagery | Motor imagery | Imagine holding a full water bottle above a table |
| OO-MI-S | Object Observation Motor Imagery Screen | Combined observation and motor imagery | Imagine holding an empty bottle of water over a table while observing the full bottle on the screen |
| OO-MI-VR | Object Observation Motor Imagery Virtual Reality | Combined observation and motor imagery | Imagine holding an empty water bottle above a table while looking at the full bottle through the VR headset |

The four experimental conditions are summarized in Table 1. They comprised two Observation Object (OO) in which participants observed a water bottle on a table on the Screen (OO–S) or through the Virtual Reality headset (OO–VR), and two combined Object Observation–Motor Imagery (OO–MI) in which participants imagined holding a 1.5L full of water bottle (without label) above a table while observing the bottle on the table in a neutral environment (Figure 1.A), either on the screen (OO–MI–S) or through the VR headset (OO–MI–VR). The virtual environment and the water bottle (Figure 1.A.b) were created by the authors using the software Unity 2023.2.9. (Unity Technologies, San Francisco, CA, USA), and a High Dynamic Range Processing (HDRP) pipeline to enhance signal robustness. The two control conditions were Rest (R) and Motor Imagery (MI). In R, participants fixated on a cross at the center of the screen. In MI, they imagined holding a 1.5L water bottle (without any label) above a table while fixating on the cross. In all conditions involving MI, participants were instructed to focus on kinesthetic imagery, given its greater effect on corticospinal excitability compared to visual imagery (24), and they had to rate the quality of their imagery on a Likert scale from 1 (poor) to 7 (excellent). In all experimental conditions, participants assessed their general fatigue using the French version of the rating-of-fatigue scale from 0 (not tired at all) to 10 (total fatigue; Brownstein et al., 2021).

The R condition was always performed first, followed by the two OO conditions. Then, the MI condition was performed, followed by the two combined OO–MI conditions. This order ensured that MI was performed after OO and before OO–MI, preventing participants from engaging in combined imagery and observation during OO and MI trials, or in motor imagery during R trials. Only the OO and OO–MI conditions were pseudo-randomized across participants (Figure 1.B).

### Questionnaire measures

Participants completed the Movement Imagery Questionnaire-third version (MIQ; French version: (61) to assess MI ability of participants in kinesthetic, internal and external visual forms of MI. After the experiment, participants completed the French version of the Simulator Sickness Questionnaire to verify the absence of VR-induced discomfort and sickness symptoms (62,63).

### Electromyographic recordings

Electromyographic (EMG) recordings were collected on the right forearm from the agonist muscles APB and PL, involved in the imagined movement, as well as the antagonist muscle ECR. Electrodes were positioned over the muscle bellies, on the thenar eminence for the APB, at one-third of the distance from the medial epicondyle and the radial styloid for PL muscle and at one third of the distance from the lateral epicondyle and the radial styloid for ECR muscle. The reference electrode was placed on the epicondyle of left elbow. Prior to the placement of two silver chloride surface electrodes (diameter: 8 mm; center-to-center distance: 2 cm; Contrôle Graphique S.A., Brie-Comte-Robert, France), the skin was shaved and cleaned with alcohol to reduce impedance to below 5kΩ. The EMG signals were amplified (gain = 1000) and filtered within a bandwidth of 100 Hz to 2 kHz, then sampled at 4 kHz using the Powerlab acquisition system (Powerlab 16/35, AdInstruments, Sydney, Australia).

### Transcranial magnetic stimulation (TMS)

TMS was applied with a 70 mm-diameter figure-eight coil connected to a Magstim magnetic stimulator (Magstim Company Ltd, Great Britain) which delivered biphasic pulses. The coil was positioned over the left motor area in a posteroanterior orientation to provide optimal results as mentioned in Loporto et al. (2011). The hotspot was identified as the scalp site which produced MEPs with the highest peak-to-peak amplitudes from the right APB muscle, using a stimulation of 60% maximum stimulator output (e.g., Wright et al., 2014). The hotspot was marked on a swimming cap participants wore during the entire session, to ensure a constant coil positioning throughout the experiment. The resting motor threshold (RMT) was then determined for each participant to be the lowest stimulation intensity that induces MEPs of 50 *µV* peak-to-peak amplitude in 50% of 10 consecutive trials while the muscle was at rest (Rossini et al., 1994). During the experiment, stimulation intensity was set to 120% and 150% of RMT. TMS pulses were delivered at variable time intervals after trial onset to prevent anticipation of the stimulation.

### Median nerve stimulation: maximal M-wave measure

Single rectangular pulses (1 ms duration) were applied to the median nerve using a constant-current stimulator (Digitimer DS7, Hertfordshire, UK) to elicit M-waves in the APB muscle. A surface bar electrode (ADInstruments, Sydney, Australia) was positioned on the distal anterior forearm, 1 cm lateral to the carpal tunnel. After identifying the stimulation site that produced the largest peak-to-peak response, the electrode was secured to maintain response consistency. Stimulation began at a low intensity (1 mA), which was then gradually increased in steps of 0.5 mA until the maximal M-wave was obtained. The intensity was subsequently increased by 20% to ensure that the M-wave remained at the plateau of its maximal amplitude. This supramaximal intensity was used to record the maximal M-wave (Mmax), thereby minimizing any influence of axonal excitability fluctuations on M-wave amplitude (64). This stimulation intensity was used throughout the experiment, for experimental and control conditions, to normalize MEPs.

### Data analysis

For each trial, peak-to-peak motor-evoked potential (MEP) amplitudes were measured in the abductor pollicis brevis (APB), palmaris longus (PL), and extensor carpi radialis (ECR) muscles. Peak-to-peak M-wave amplitudes were also measured in the APB muscle. For each participant, condition, and stimulation intensity, mean amplitudes were then calculated separately for each muscle. To account for peripheral excitability, APB MEP amplitudes were normalized to the maximal M-wave amplitude (MEP/Mmax ratio). Because M-waves were recorded only from APB muscle, PL and ECR MEP amplitudes were analyzed as absolute values.

All statistical analyses were performed using R (version 4.4.3; R Core Team, 2025). Analyses were conducted separately for each muscle and stimulation intensity. The effect of Condition (R, OO–S, OO– VR, MI, OO–MI–S, and OO–MI–VR) on MEP amplitude was tested using linear mixed-effects models, with Condition as a fixed effect and Participant as a random intercept to account for repeated measures within participants. To check for normality of residuals and homoscedasticity, both were carefully examined using quantile–quantile plots and residual plots, respectively.

The significance of the fixed effect of Condition was assessed using Type III ANOVA. When a significant main effect was observed, pairwise comparisons were performed with Bonferroni-corrected *p*-values. In addition, a priori planned contrasts were used to test the specific effects of motor imagery, affordances, and display modality, as detailed in Table 2. Effect sizes were reported as partial eta squared 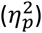 for the ANOVA and as Cohen’s *d* for pairwise and planned comparisons. The alpha level was set to 0.05.

**Table 2.**
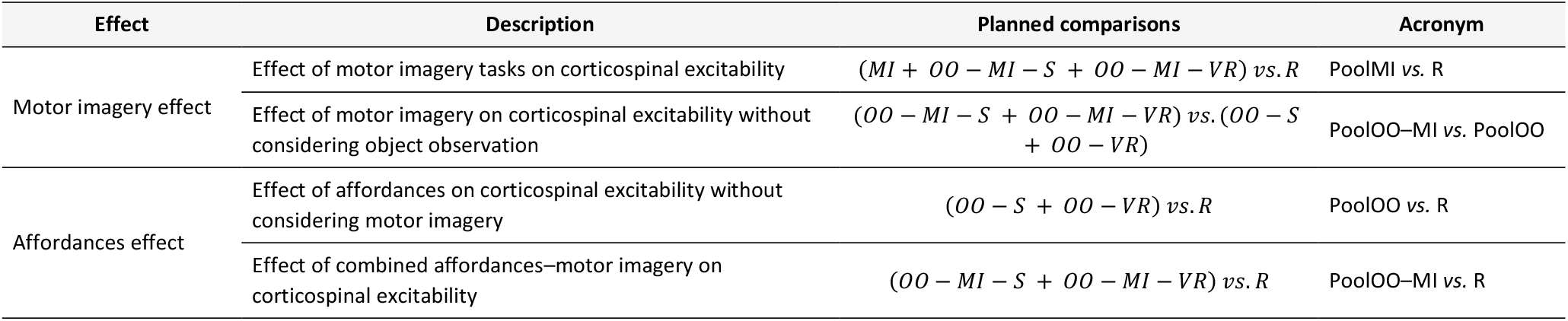
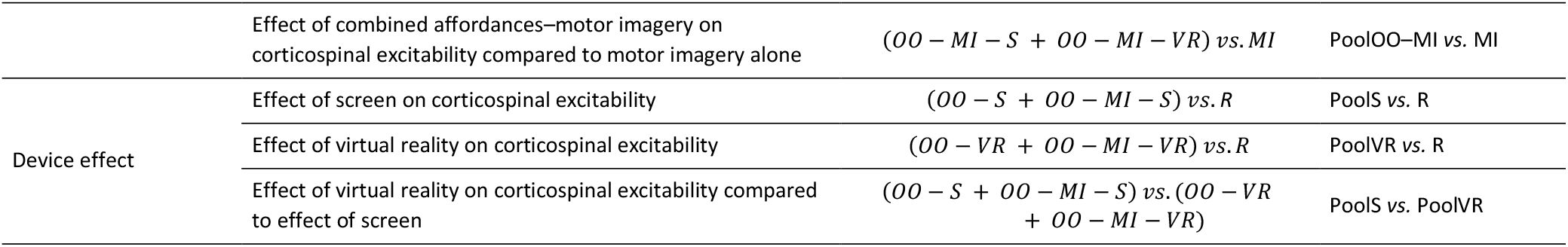
Description of planned comparisons for Motor imagery effect, Affordances effect, and Device effect. MI: Motor Imagery; OO: Object Observation; R: Rest; S: Screen; VR: Virtual Reality.

Pre-stimulus EMG activity was quantified as the root mean square (RMS) of the signal during the 100 ms preceding each TMS pulse. To verify the absence of relevant background muscle activity, trials were visually inspected, and only trials with pre-stimulus EMG RMS below an a priori resting-EMG threshold of 0.02 mV were retained for analysis (66).

Fatigue ratings across the six conditions and and motor imagery quality across the three motor imagery conditions (MI, OO–MI–S, and OO–MI–VR) were analyzed using repeated-measures ANOVA when normality was met, or a Friedman test otherwise. When a significant main effect was observed, post-hoc pairwise comparisons were conducted using paired *t* tests or Wilcoxon signed-rank tests, with Bonferroni correction for multiple comparisons. Effect sizes were reported as Cohen’s *d* for *t* tests and rank-biserial correlations for Wilcoxon tests.

## RESULTS

### Fatigue and quality of motor imagery

No significant differences were observed across conditions for either fatigue [Friedman test: χ^2^(5) = 10.07, *p* = .073, Kendall’s *W* = .168] or motor imagery quality [repeated-measures ANOVA: F(2, 22) = 1.157, *p* = .333, 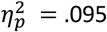].

### Motor-evoked potentials

The mean RMT was 54.40% of maximal stimulator output (±*SD* = 7.95).

For the 120% stimulation intensity (Figure 2.A.), no significant differences were observed across conditions for MEPs recorded from APB, ECR, and PL (all *ps* > .10).

**Figure 2.**
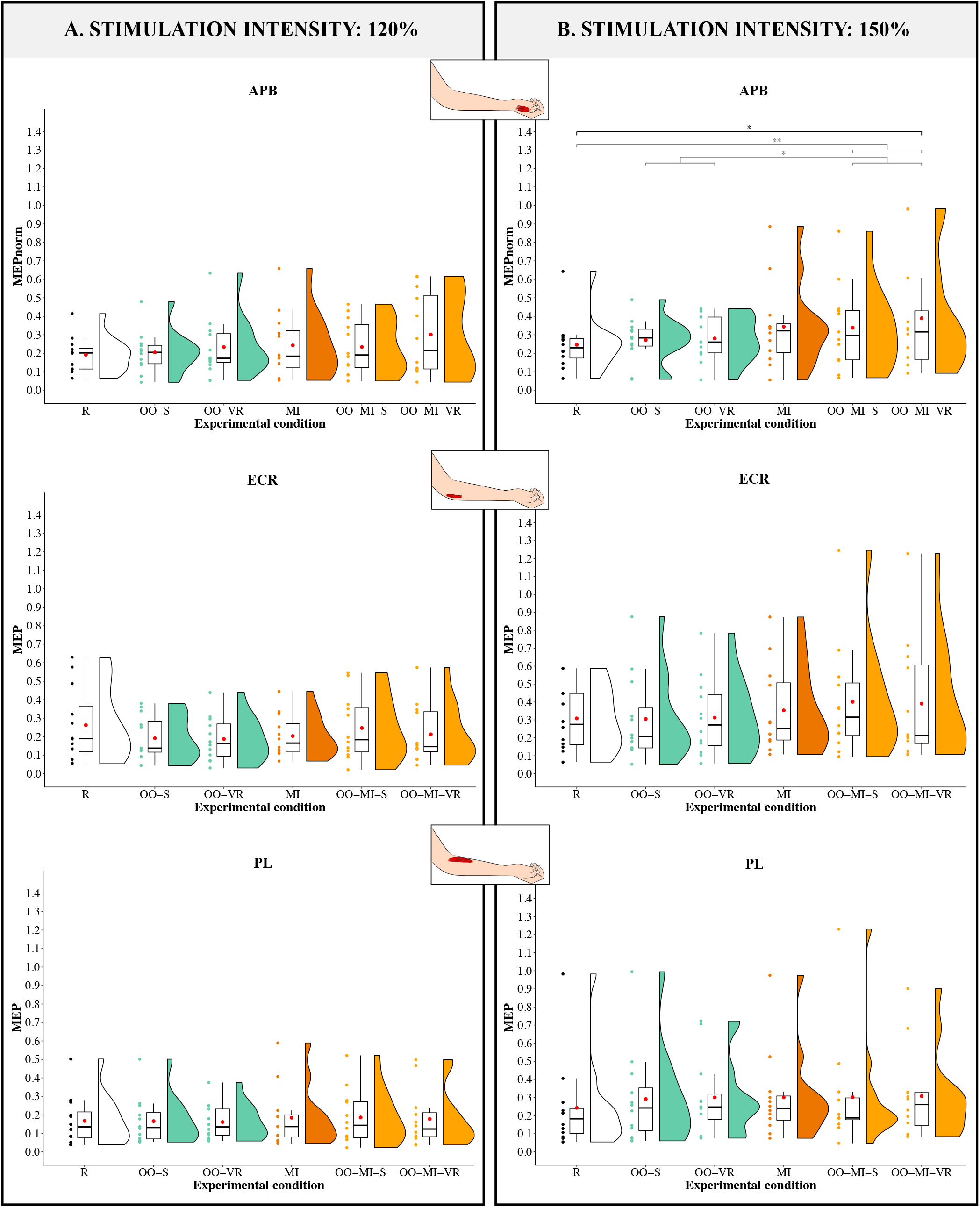
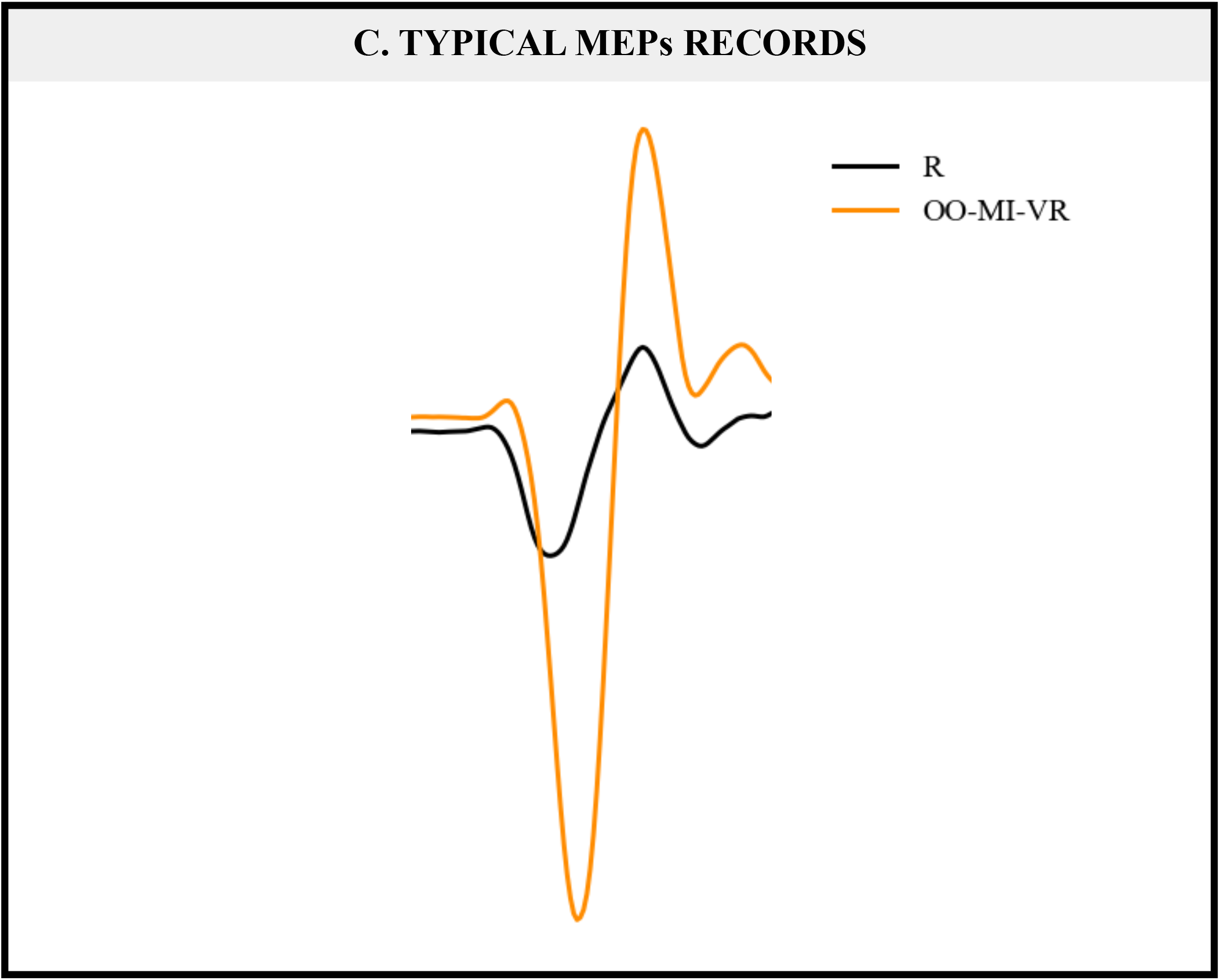
MEPs obtained from Abductor Pollicis Brevis (APB), Extensor Carpi Radialis (ECR), and Palmaris Longus (PL) muscles for each stimulation intensity. MEP: Motor evoked potentials; normMEP: normalized MEP corresponding to MEP/M-wave ratio; R: Rest; OO–S: Object Observation Screen; OO– VR: Object Observation Virtual Reality; MI: Motor Imagery; OO–MI–S: Object Observation Motor Imagery Screen; OO–MI–VR: Object Observation Motor Imagery Virtual Reality. The main effect is symbolized in dark, planned comparisons are represented in gray; *: p < 0.05; **: p < 0.01; **A**. Stimulation intensity: 120%; **B**. Stimulation intensity: 150%. **C**. Typical MEPs records obtained for a participant during Rest (R; black line) and Object Observation Motor Imagery Virtual Reality (OO-MI-VR; orange line), for the 150% stimulation intensity.

For the 150% stimulation intensity (Figure 2.B.), a significant main effect of *Condition* was observed for MEPs recorded from APB [ANOVA: *F*(5,55) = 3.063, *p* = .016; 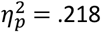]. Bonferroni-corrected post hoc showed significantly larger MEP amplitudes in OO–MI–VR than in R (*p* = .027; *d* = 1.338). Figure 2.C. shows typical MEP traces for these two conditions, highlighting the differences between them. Planned comparisons indicated that corticospinal excitability significantly increased when affordances were combined with motor imagery [(OO–MI–S + OO–MI–VR) *vs*. R; *p* = .009; *d* = 1.097], and during motor imagery without considering affordances [(OO–MI–S + OO–MI–VR) *vs*. (OO–S + OO–VR); *p* = .013; *d* = 0.819]. Only the use of VR to display the object tended to increase corticospinal excitability [(OO–VR + OO–MI–VR) vs. R; *p* = .068; *d* = .829]. No significant changes were observed for MEPs from ECR and PL (all *ps* > .50).

## DISCUSSION

### Enhancement of corticospinal excitability with combined affordances and motor imagery

Consistent with the notion that affordances alone increase corticospinal excitability, our study showed that this facilitation is further enhanced when OO is combined with MI of an action involving that object. It is now established that combined AO+MI engages motor-related brain areas and facilitates corticospinal excitability more than AO or MI alone (30,31,36,38,39). In affordances-based contexts, Bordoloi et al. (2022) recently reported an increased N2 amplitude—a marker of affordance detection and motor preparation—during combined AO+MI relative to either AO or MI alone, suggesting that perceiving affordances leads to a clearer mental simulation of the associated action (67). Our findings extend this idea by providing the first demonstration of increased corticospinal excitability during OO+MI compared to rest or OO alone. This indicates that visual presentation of an object can prime motor actions even when the action is predetermined through MI. The motor system can use information from object features to select the appropriate motor schema within the motor repertoire (68), with objects providing a clear target for imagined actions, such as grasping a bottle. Premotor and primary motor areas are recruited during actual grasping movements, reflecting visuospatial processing and coordination of hand movements (69,70). Visuospatial attention would be naturally directed towards objects that afford action, optimizing the coupling between visual attention and motor affordance (71). Schulz and collaborators (2018) showed an increased activation of the supplementary motor area (SMA) during MI of grasping an everyday object (i.e., a bottle) compared with a geometrical shape (i.e., a cone), consistent with the notion that object perception automatically triggers action affordances (48). The SMA plays a key role in movement selection and inhibitory control, preventing involuntary execution of motor scripts evoked by object affordances (72–74). Taken together, these mechanisms suggest that object perception automatically activates object-related motor programs, thereby potentiating associated motor actions (52,75). This interpretation fits with the idea that our sensorimotor system transforms visual perception of objects features into action-relevant coordinates before selecting the most appropriate motor response (76). A next step for future research would be to manipulate the properties of the object to determine whether corticospinal excitability scales in a comparable manner across different affordance configurations.

### Immersive object observation and motor imagery, the ideal combination to increase corticospinal excitability?

Our main finding is that combining MI with object observation in virtual reality significantly facilitates corticospinal excitability—a result that, to our knowledge, has not been previously demonstrated. VR headsets create an illusion of reality by blurring the boundary between the real and virtual worlds. High-quality graphics and multisensory feedback allow users to experience immersion, which is strengthened by the sense of presence achieved through affordances and embodiment (42,77,78). Compared with conventional screen-based displays, VR environments facilitate the recruitment of motor circuits, especially when a manipulable object is visible, likely due to improved spatial discrimination and enhanced rhythmic motor patterns (42). Previous studies have shown that VR-based action observation can facilitate MI (79–81), likely due to enhanced cortical neuroplasticity compared to screen-based observation. This is reflected by a larger cerebral network activation than MI alone, involving premotor, motor, parietal and occipital areas (43). VR environments have been shown to simulate real-world actions and provide multisensory feedback, resulting in greater improvements in manual dexterity in healthy participants (82) and shorter rehabilitation time with better recovery in stroke patients (83–85). Neurophysiological studies showed a larger MI-induced event-related desynchronisation, reflecting increased cortical activation, in the contralateral motor cortex, when using VR compared to any visual presentation (44), only during but not before the MI practice (42). In our study, combining MI with congruent VR-based OO may facilitate corticospinal excitability at a less time cost compared to no visual presentation or screen presentation. Moreover, using an object instead of an action during observation alleviates the constraints of temporal and coordination congruency required when the imagined and observed actions must match (86,87). Our results are promising for rehabilitation and physical training, as the realistic visual representation and spatial-sensory grounding of the object generate more engaging affordances that activate the motor system. We can assume that this facilitation observed with combined OO–MI–VR may result from an interaction between immersive affordance and MI, although further studies are needed to clarify this mechanism.

### Changes depend on muscle and stimulation intensity

The observed facilitation of corticospinal excitability is muscle-dependent. First, MI modulates corticospinal excitability in a muscle-specific manner, targeting to the agonist muscle of the imagined action (10,88). This specificity is well established, as MI-related activation of motor regions is highly movement-dependent, event for close muscles—for example, elbow-flexion MI increases corticospinal excitability in the biceps but not in the triceps(10). During object observation, visual priming is also muscle-specific with modulation of excitability restricted to the cortical representations of agonist muscles (89,90). For example, observing pinchable objects has been shown to increase corticospinal excitability in the FDI but not in the APB (75). Second, the effect is muscle-specific regarding the area with a modulation in the thumb (APB) rather than the forearm (PL). This could stem from the fact that 1) the APB has a larger cortical representation, as a hand muscle, than PL, making it more sensitive to MI- and VR-related interventions as more pyramidal neurons are available and/or 2) participants may focus more naturally on hand than forearm muscles while imagining movement as grasping, leading to stronger APB activating. This discrepancy might also reflect a methodological bias, since TMS parameters were optimized for APB rather than other muscles. Consequently, the hotspot and stimulation threshold intensity may not be the best one to obtain the optimal MEP in PL and ECR. However, given the spatial accuracy of TMS and the proximity of their motor areas, it can be assumed that a reliable MEP can still be obtained together with the same hotspot.

An important methodological consideration arises from our study, indicating that the most suitable stimulation intensity for detecting changes was 150% of RMT. No significant modulation was observed at 120% of RMT. It has already been shown that this classical stimulation intensity is insufficient to detect differences between MI and rest or minimal voluntary contraction (25). Therefore, higher intensities may be more appropriate for detecting even slighter changes between different cognitive tasks.

### Conclusion

A central question in the MI literature is how to identify the optimal combination of conditions to maximize its benefits. Numerous studies have investigated the combination of MI and action observation, given its established role in facilitating corticospinal excitability. However, some limitations emerged, particularly regarding the temporal congruence between observed and imagined actions, as well as the sense of ownership or identification with the observed limbs or performer (human or avatar). The present study demonstrates a significant facilitation of corticospinal excitability when MI is combined with object observation in a virtual environment. This modulation provides the first evidence for enhanced MI through VR-based affordances perception. We suggest that this effect may result from stronger activation within motor-related areas when affordances are coupled with MI, thereby potentiating the benefits of MI. These findings offer new insights into MI-induced corticospinal excitability and may have potential applications in rehabilitation and sports training. Further studies are needed to explore the optimal parameters for this combined intervention and identify neurophysiological markers of cortical modulation.

## DATA AVAILABILITY

Source data for this study are openly available at https://doi.org/10.17605/OSF.IO/UNG7H

## ACKNOWLEDGMENTS

The authors thank all the participants who engaged in this study.

## AUTHOR CONTRIBUTIONS

C.B., S.A., and S.G. conceived and designed research; C.B. and M.B. performed experiments; C.B. analyzed data; C.B. and S.G. interpreted results of experiments; C.B. prepared figures; C.B. drafted manuscript; C.B. and S.G. edited and revised manuscript. C.B., M.B., S.A., and S.G. approved final version of manuscript.

## TABLES

Tables must be submitted in Word table format, not as tabbed text or as embedded images. Do not include shading, color highlights, or embedded graphics. Notes, statistical measures of variations (SD, SE and others symbols), and abbreviations should be defined in the table legend. Follow <u>Rigor and Reproducibility Guidelines (PDF)</u> for more guidance. Example:

